# Two ecotype-related long non-coding RNAs in the environmental control of root growth

**DOI:** 10.1101/579656

**Authors:** Thomas Blein, Coline Balzergue, Thomas Roulé, Marc Gabriel, Laetitia Scalisi, Céline Sorin, Aurélie Christ, Etienne Delannoy, Marie-Laure Martin-Magniette, Laurent Nussaume, Caroline Hartmann, Daniel Gautheret, Thierry Desnos, Martin Crespi

**Affiliations:** Institute of Plant Sciences Paris Saclay IPS2, CNRS, INRA, Université Paris-Sud, Université Evry, Université Paris-Saclay, Bâtiment 630, 91405 Orsay, France; Institute of Plant Sciences Paris-Saclay IPS2, Paris Diderot, Sorbonne Paris-Cité, Bâtiment 630, 91405, Orsay, France.; CEA, Institut de Biologie Environnementale et de Biotechnologie, Laboratoire de Biologie du Développement des Plantes, Saint-Paul-lez-Durance, F-13108, France.; CNRS, Unité Mixte de Recherche 7265 Biologie Végétale & Microbiologie Environnementale, Saint-Paul-lez-Durance, F-13108, France.; Aix-Marseille Université, Saint-Paul-lez-Durance, F-13108, France.; UMR MIA-Paris, AgroParisTech, INRA, Université Paris-Saclay, 75005, Paris, France; I2BC, Institute for Integrative Biology of the Cell, CEA, CNRS, Université Paris Sud, 1 avenue de la terrasse, 91198 Gif sur Yvette, France.

## Abstract

**Background:** Root architecture varies widely between species and even between ecotypes of the same species despite the strong conservation of the protein-coding portion of their genomes. In contrast, non-coding RNAs evolved rapidly between ecotypes and may control their differential responses to the environment as several long non-coding RNAs (lncRNAs) can quantitatively regulate gene expression.

**Results:** Roots from Columbia (Col) and Landsberg *erecta* (L*er*) ecotypes respond differently to phosphate starvation. We compared complete transcriptomes (mRNAs, lncRNAs and small RNAs) of root tips from these two ecotypes during early phosphate starvation. We identified thousands of new lncRNAs categorized as intergenic or antisense RNAs that were largely conserved at DNA level in these ecotypes. In contrast to coding genes, many lncRNAs were specifically transcribed in one ecotype and/or differentially expressed between ecotypes independently of the phosphate condition. These ecotype-related lncRNAs were characterized by analyzing their sequence variability among plants and their link with siRNAs. Our analysis identified 675 lncRNAs differentially expressed between the two ecotypes including specific antisense RNAs targeting key regulators of root growth responses. Mis-regulation of several intergenic lncRNAs showed that at least two ecotype-related lncRNAs regulate primary root growth in Col.

**Conclusions:** The in depth exploration of the non-coding transcriptome of two ecotypes identified thousands of new lncRNAs showing specific expression in root apexes. De-regulation of two ecotype-related lncRNAs revealed a new pathway involved in the regulation of primary root growth. The non-coding genome may reveal novel mechanisms involved in ecotype adaptation of roots to different soil environments.

## Introduction

Over the last decade, genome-wide transcriptomics studies revealed that a large part of eukaryotic genomes, intergenic to protein-coding genes, was transcribed. These molecules, globally known as non-coding RNAs [1] may regulate genome expression at transcriptional, post-transcriptional and epigenetic levels and are generally divided in small (21 to 24 nt) and long (> 200nt to 100 kb) non-coding RNAs. Plant small RNAs are processed from longer non-coding transcripts that generally contain a hairpin structure or lead to double stranded RNA formation. Plant small RNAs include miRNAs, endogenous siRNAs (both generally 21-22 nt long) and the most abundant heterochromatin siRNA (hc-siRNAs with a length of 24 nt [2]). On the other hand, long non–coding RNAs (lncRNAs) are a heterogeneous group of RNA molecules with a coding capacity shorter than 50 amino acids [3,4]. LncRNA transcripts are generally polyadenylated (polyA) and can be intergenic (lincRNAs), intronic (incRNAs) or natural antisense (NATs) with respect of protein coding genes [5,6]. When compared to mRNAs, lncRNAs are expressed at low levels in a tissue-specific manner or in response to environmental stresses [7,8] and are more frequently accumulated in the nucleus relative to the cytoplasm [9]. Recently, some RNA motifs that promote nuclear retention have been identified in lncRNAs (reviewed in [10]). In human cells, Mukherjee et al [11] observed that lncRNA showed lower synthesis and higher degradation rates than protein-coding mRNAs. [11].

LncRNAs utilize both cis and *trans* modalities of action to regulate gene expression through interaction with ribonucleoproteins and can form scaffolds and/or sequester proteins or RNA molecules as decoys or sponges. However, molecular function has only been identified for a very small proportion of lncRNAs both in animals and plants. As lncRNA genes lack regions with high primary sequence constraints [9], it is difficult to use sequence conservation to identify potential functions as it has generally been done for protein coding genes or housekeeping RNA genes [12,13].

Recently, the utilization of new methods (Structure-seq [14] SHAPE-MaP [15] and icSHAPE [16]) allows the prediction of lncRNA structures *in vivo* (named RNA structuromes, [17]). It has also been shown that these RNA structures can be modified under different environmental conditions such as temperature or metabolic changes.

It is noteworthy that in 16 vertebrate species, Herzoni et al. [18] have shown that thousands of lncRNAs, without sequence conservation, appear in syntenic positions and some present conserved promoters [7,18–20]. In rice and maize, the characterization of lncRNAs also revealed a higher positional conservation (around 7 times more) than sequence conservation [21]. In the same way, in Brassicaceae, the fraction of lincRNAs, for which sequence conservations with Arabidopsis could be detected, decreased as the phylogenetic distance increased [22]. In fact, emerging mechanisms involved lncRNAs in the regulation of genome expression, however, we cannot exclude that part of lncRNAs could be simply transcriptional by-products and/or that the sole act of their transcription rather than their sequence, is the source of the regulation activity [23,24].

This last decade, whole-genome variations and evolution in many model species had been determined through resequencing approaches (from yeast and humans to Arabidopsis, rice and certain crops). These studies provided the characterization of pan-genomes composed of “core” genomes (present in all accessions) and “dispensable” genomes (specific to two or more accessions or cultivars or even unique sequences specific to only one accession). Core genes are frequently highly expressed while dispensable genes are variably expressed, and generally in a tissue specific manner [25]. The dispensable genomes may play important roles in the capacity of individual organisms to cope with environmental conditions [26]. Indeed, identification of natural variations in large worldwide populations (accessions) of *Arabidopsis* ([27] and more recently in [28]) showed an average of one SNP per 10 bp more frequently located in intergenic regions than in coding mRNAs [28]; this has also been observed recently in rice for which only 3,5 % of SNP and 2,5 % of small InDels were located in the coding region [29]. This latter observation would explain why lncRNAs differ between closely related species (e.g. rat and mouse, [30], human and chimp, [31] or different plant species, [32]). Recently, lncRNAs were found to vary, at the individual level, much more than the protein coding genes [33]. These modifications may drive new regulatory patterns of gene expression as lncRNA evolved quickly and many of them emerged recently [31]. Indeed, regulations derived from novel transcription units due to transposon rearrangements were suggested to be highly variable until they are selected for a strict regulatory pattern [34].

The Col and L*er* accessions display different primary root growth and architecture in their response to phosphate starvation [35]. The identification of LPR1, a major QTL, has been done in RIL lines obtained by the crosses of accessions presenting this opposite root response to low Pi [35,36]. Pi deficiency is perceived at the root apex [36]. When the primary root tip of a Col seedling encounters a low Pi medium, cell elongation in the elongation zone rapidly decreases and cell proliferation in the RAM progressively ceases [37–40]. By contrast, in L*er* seedlings elongation and proliferation of root cells continue, thereby sustaining root growth. In addition, the root tip concentrate a high proportion of Pi transporters which provide an important contribution to Pi nutrition and Pi systemic signaling [41,42]. Hence, we decided to identify and characterize the non-coding transcriptomes of Col and L*er* root apexes during early phosphate starvation responses to search for lncRNAs potentially linked to this differential growth response. Interestingly, we identified thousands of new Arabidopsis lncRNAs, notably in the L*er* accession, with only a minor fraction being linked to small RNA production. Several “ecotype-specific” or “-enriched” variants, highly conserved at DNA level, showed expression variation correlated with changes in the expression of key regulators of phosphate starvation response. Functional analysis of 6 lncRNAs in Col revealed two new regulators of primary root growth allowing us to hypothesize that novel lncRNA expression patterns contribute to the modulation of environmental responses in different ecotypes.

## Results

### Columbia and Landsberg root tip transcriptome assemblies

We first reconstructed the transcriptome of root tip of Columbia (Col) and Landsberg *erecta* (L*er*) ecotypes. These two ecotypes present contrasting root phenotypes in response to Pi deficiency [35]. The root growth arrest of the Col ecotype occurred in the first hours of low phosphate sensing by the root tip [38]. Therefore, in these two ecotypes, we performed comparative whole genome transcriptomic analyses using paired end sequencing of three biological replicates of root tips during a short kinetics (0 h, 1 h and 2 h) of low phosphate treatment (10µM, see the “Methods” section). To discard possible differences related to the erecta mutation known to be present in L*er* ecotype, we used Col^er105^ mutant carrying this mutation in Col. We obtained between 47M and 65M of reads (Additional file 1: Table S1). For each ecotype, the reads were independently mapped to their reference genome (Additional file 2: Fig. S1a): TAIR10 for Col [43] and L*er* v7 for L*er* [44]. We selected these two genome versions since they shared the same gene annotation (TAIR10). For these two genomes, we predicted new transcripts by comparing our data to those available in TAIR10 annotations. The homology of newly predicted transcripts between the Col and L*er* genomes was determined by mapping them on the other genome (TAIR10 and L*er* v7). When the newly transcripts overlapped with pre-existing annotations, fusions between the two transcripts were done to generate the new transcripts. We retained as new transcripts only RNA molecules of at least 200 nt. The L*er* v8 version of the L*er* genome lacks the official TAIR10 annotation[45]. Therefore, for the transcripts mapping on L*er* genome, we retained only the ones that also mapped on this version. Newly identified transcripts by this pipeline (Additional file 2: Fig. S1a) were then compared with the ones already described in different Arabidopsis databases: Araport 11 [46], RepTas [8], CANTATAdb [47], miRBase v21 [48] and with the results of two previous studies concerning lncRNAs [49,50]. Finally, we used the COME software [51] to determine the potential coding capacity of new identified transcripts (Additional file 2: Fig. S1a). On the basis of both database information and COME predictions we classified the corresponding genes as coding or non-coding.

In total, we identified 5313 and 6408 novel putative genes respectively in Col and L*er* ecotypes (Fig. 1a; Additional file 1: Table S2; Additional file 3). In root apexes, newly discovered genes were predominantly non-coding RNAs: 76% for Col and 77% for L*er* of total number of new genes (Fig. 1a; Additional file 2: Fig. S1b, c). This suggests that the coding capacity of the Arabidopsis genome is now well documented in TAIR10 and Araport11 databases (Additional file 2: Fig. S1b). As expected, non-coding genes were globally less expressed than coding genes (Additional file 2: Fig. S2a, b). Genes specifically detected only in one ecotype belong much more to the non-coding class (>40%) than to the coding one (<8%; Fig. 1b, c). Moreover, the proportion of ecotype-specific expressed lncRNAs was higher for intergenic lncRNA genes (52%) than for NATs (34%; Fig. 1d,e). Overall, these results show that at the ecotype level the expression of non-coding genes is more ecotype specific than for coding genes.

**Fig. 1.**
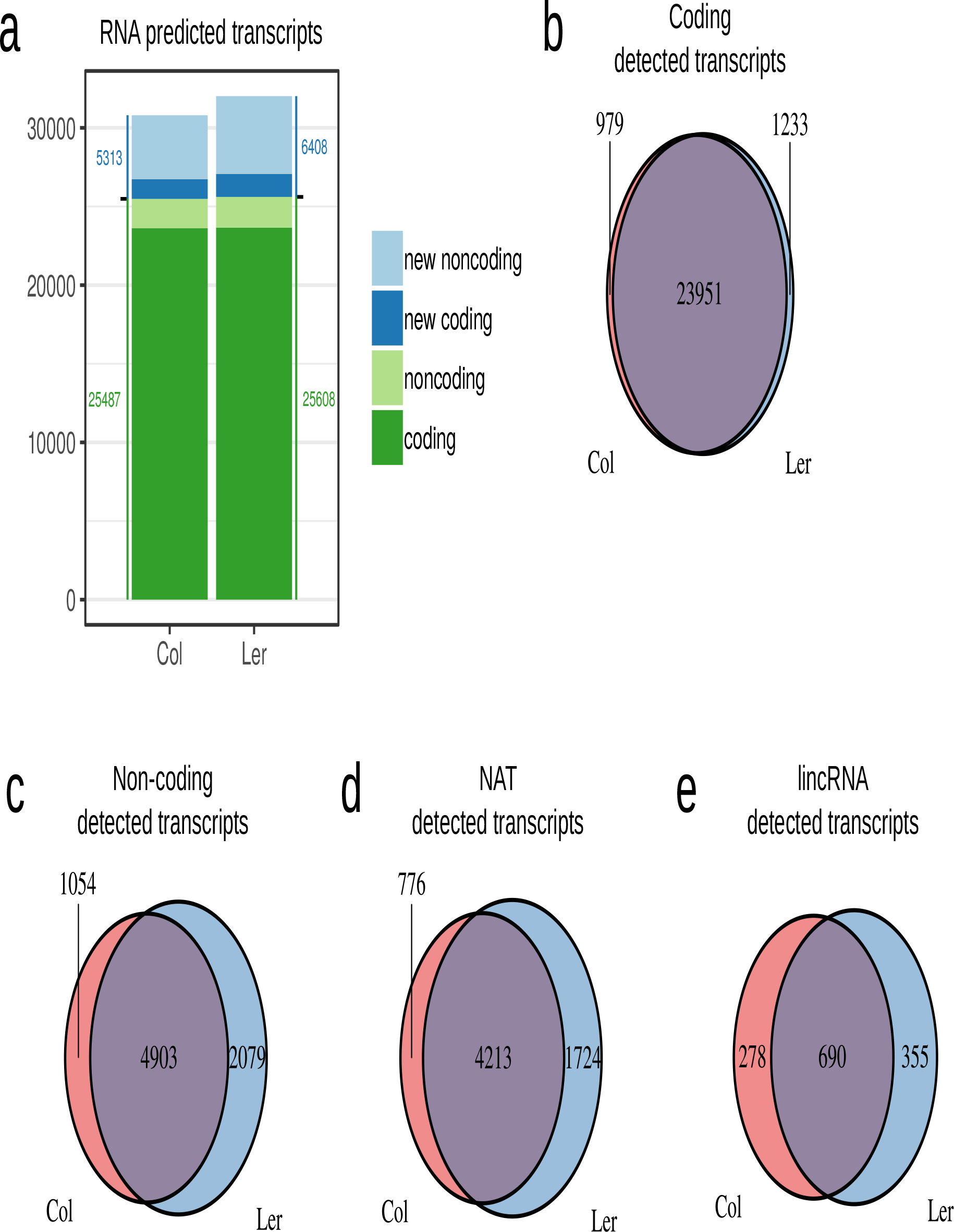
Identification of the transcripts and their repartition between the two ecotypes. (a) Number of predicted coding and non-coding transcripts in the two ecotypes classed by type. New transcripts were not discovered in previously published studies. Then, detection at RNA levels of all transcripts predicted in each ecotype: classified as coding (b) or as non-coding (c). For the latter two classes are defined if they are antisens of another annotation (NAT, d) or are intergenic (lincRNA, e). In contrast to coding genes, many ncRNAs, notably those intergenic, were detected only in one ecotype despite the DNA sequence similarity in both ecotypes.

We detected a greater number of new genes (essentially non-coding) in the L*er* ecotype (Fig. 1a, c). We wondered whether this difference might be due to a technical bias. Hence, we examined the sequencing saturation of the different libraries (see the “Methods” section; Additional file 2: Fig. S2c). We noted that in the last 2% of sequencing reads, less than 10 additional genes were newly detected. We considered then that sequencing was deep enough to detect the very large majority of expressed genes in both ecotypes. Hence, the difference of gene detection between Col and L*er* does not result from a sequencing bias.

We next sought whether these newly detected expressed L*er* genes (coding or non-coding) corresponded to specific parts of L*er* genome that were missed or rearranged in Col genome and can easily explain expression differences between ecotypes. Out of the 7357 newly identified genes, only 41 and 53 genes, respectively in Col and L*er*, coincided with missing DNA sequences in the other ecotype (Fig. 2a) showing that the DNA sequence of the different new genes is largely conserved. Thus, the large majority of the differences in transcript accumulation between ecotypes came from a shift in transcription in one of the two ecotypes of an almost identical DNA region (except for few SNPs). This change in expression could be due to the deregulation of some master gene regulator, or to the accumulation of small sequence differences in gene promoter regions with functional consequences at transcription initiation level or the consequence of specific differences in epigenetic status in the lncRNA-producing region due to TE insertions or other rearrangements at large distance from the loci.

**Fig. 2.**
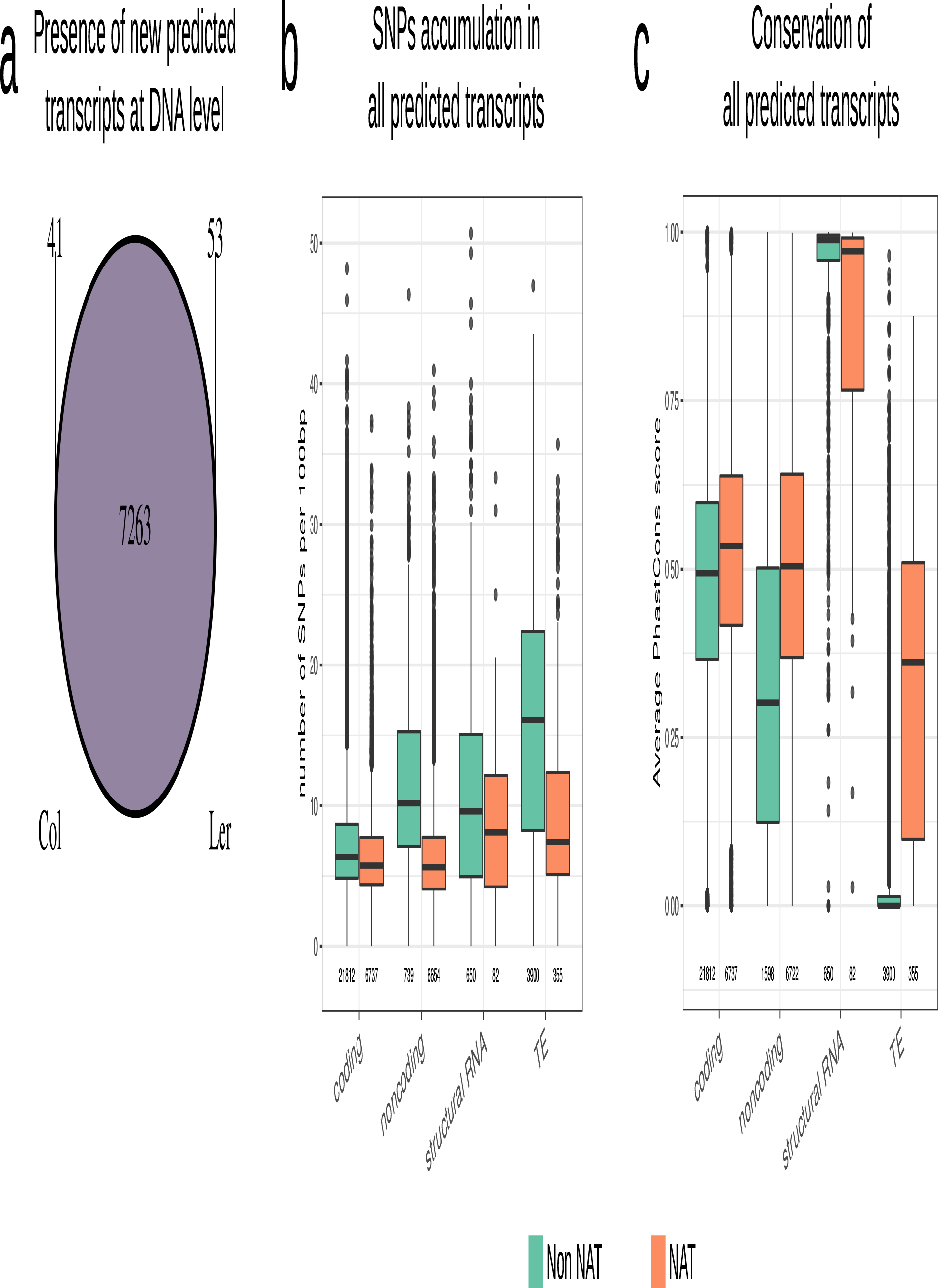
Characterization of transcript at the DNA levels (a) Detection of the DNA sequence of the newly predicted transcripts in the two ecotypes (minimum of 90% of sequence identity and minimum of 90% of RNA length). The large majority of RNAs come from common DNA regions from both ecotypes. (b) SNPs accumulation per 100 bp of transcript length for each type of transcript according to data of 1001 genomes project. (c) Conservation among plant species (average PhasCons score) of each type of transcript according to their genomic position in relation to other annotations. In (b) and (c), NAT are natural antisense RNAs (overlapping with another mRNA, see Materials and Methods) and Non-NAT are transcript without any transcript overlapping on the other DNA strand (intergenic).

### Evolutionary analysis of lncRNA genes expressed in root tips

We then characterized the evolution of Arabidopsis genes expressed in root tips profiting from the extensive sequence information in Arabidopsis accessions [28]. According to current annotations, genes were classified as non-NAT (no gene on the other strand) or NAT (presence of a gene on the other strand). First at the level of the *Arabidopsis thaliana* species, we calculated the rate of SNP accumulated in each gene among all the different ecotypes (Fig. 2b). As expected, transposable elements (TE) accumulated much more SNPs than coding genes, whereas non-NAT lncRNAs and structural RNA genes showed an intermediate level of SNPs between TEs and coding genes. By contrast, the level of SNPs was generally similar for NAT lncRNA and coding genes as the major part of NAT lncRNA genes are antisense of coding genes which are under clear selection pressure.

In a second step, to investigate sequence evolution at a larger scale, we used the PhastCons score that represents an inter-species level of nucleotide conservation (normalized between 0 and 1) according to the alignment of 20 Angiosperm genomes (Fig. 2c, [52]). As expected, structural RNA genes were strongly conserved (median PhastCons score of 1), while transposable elements were not (median PhastCons score of 0). Coding genes present a score between these two extremes (around 0.5). Interestingly, non-NAT lncRNA genes showed an intermediate score between coding and transposable genes (median PhastCons score around 0.3) whereas NAT lncRNA genes had again the same degree of conservation than coding genes. These observations further suggest that the sequence of the NAT lncRNA genes are strongly constrained, likely due to the strong selection pressure on their overlapping coding genes, whereas intergenic non-coding genes allow more variability even though they are more constrained than TEs.

### Few lncRNA transcripts co-localize with small RNA generating loci

In animals and plants, some lncRNA loci co-localize with genomic regions producing small RNA molecules [53,54]. Therefore, we asked whether lncRNAs loci expressed in root tips could generate small RNAs. Small RNA sequencing was done on similar samples used for the long RNA sequencing and the identification of small RNAs was conducted independently on each genome (TAIR10 and L*er* v7; see the “Methods” section). Then, small RNAs were mapped on genes. The majority of the lncRNAs containing small RNAs generated non-phased molecules of 21/22nt or 24nt (Additional file 2: Fig. S3a, b). A minor proportion of lncRNAs overlaped with phasiRNAs or were likely miRNA precursors (Additional file 2: Fig. S3a, b).

We then analyzed the potential link between small RNAs and lncRNAs in each ecotype. The majority of lncRNAs did not accumulate siRNAs (6452 genes out of 7850 detected lncRNAs) that were specifically observed in one ecotype. Many of them also did not generate any siRNA on either ecotype (2688 genes out of 3110 ecotype specific detected genes; Fig. 3a, long in Col and ND in L*er* or *vice versa*). Thus, the differential presence of lncRNA between ecotypes could not be linked to a change in siRNA production from the encoding loci.

**Fig. 3.**
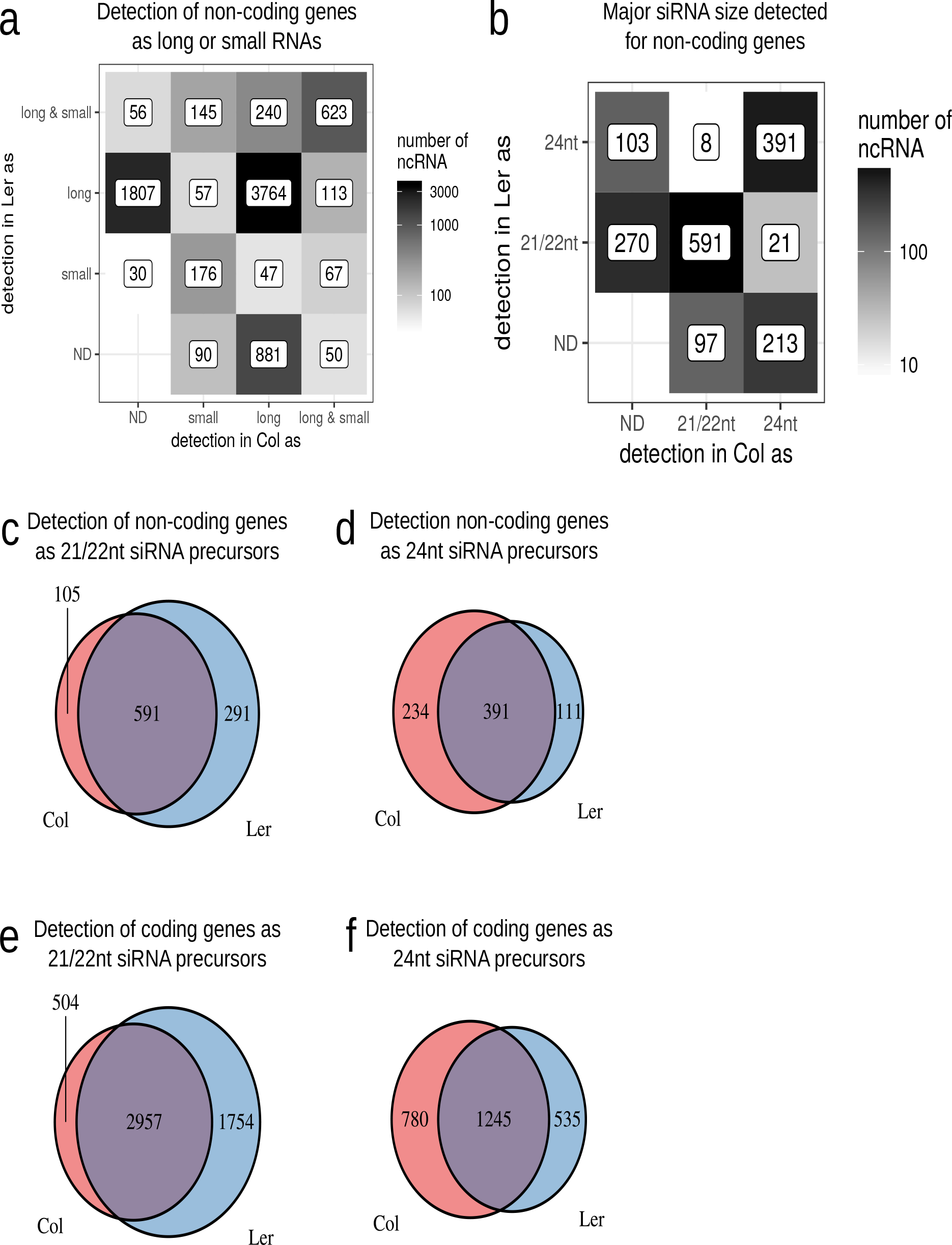
Long ncRNAs as small RNA precursors (a) Detection specificity of non-coding transcript as siRNAs (at the level of 1 RPM) or long RNA (at the level of 1 TPM) between the two accessions. ND = not detected. The major difference between Col and L*er* is the lncRNA component of the transcriptome. (b) Repartition of the major siRNA size for non-coding transcripts detected as long and short (level of 1 RPM) in the two ecotypes. ND = not detected. There is no major change of siRNA size between the two accessions. (c) Detection specificity of non-coding RNA as 21 nt and 22 nt siRNAs precursors at the level of 1 RPM in each ecotype. (d) Detection specificity of non-coding RNA as 24 nt siRNAs precursors at the level of 1 RPM in each ecotype. (e) Detection specificity of coding RNA as 21 nt and 22 nt siRNAs precursors at the level of 1 RPM in each ecotype. (f) Detection specificity of coding RNA as 24 nt siRNAs precursors at the level of 1 RPM in each ecotype.

We then wondered whether different small RNA processing from lncRNAs could occur between the two ecotypes. We first looked at lncRNA that accumulated small RNA only in one accession: (i) lncRNAs that could generate siRNAs in only one ecotype while detected as lncRNAs in both ecotypes (long in L*er* and long plus small in Col and vice versa, respectively 113 and 240 genes) and (ii) loci that produced only siRNAs in one ecotype and only detected as lncRNAs in the other one (long in L*er* and small in Col or vice versa, respectively 57 and 47 genes). It is known that 21/22nt siRNAs act on gene transcripts whereas 24nt siRNAs mediate chromatin modifications [53,54]. Thus, alterations in the size of specific siRNAs, in one ecotype, could indicate a modification of the type of gene regulation: post-transcriptional (21/22nt) or epigenetic (24nt) in the other ecotype. Therefore, we analyzed the major small RNA size that accumulated on lncRNA genes. Among the lncRNA genes accumulating siRNA, a large portion accumulated the same size in both ecotypes (591 for 21/22nt siRNA and 391 for 24nt siRNA) or produced siRNA in one ecotype (367 for 21/22nt and 316 for 24nt, Fig. 3b). Only 29 lncRNA genes accumulated a different size of siRNAs between the two ecotypes among the 1694 lncRNA genes accumulating siRNAs. Therefore, no major change of reciprocal posttranscriptional or transcriptional regulation of lncRNA by small RNAs could be established between ecotypes.

We finally investigated the specificity of detection of siRNAs between the two ecotypes. First, we studied lncRNA genes predicted to produce phased 21/22nt siRNAs. Among the 7 predicted lncRNAs, only 2 were specific to L*er* (Additional file 2: Fig. S3d). Second, we searched for miRNA loci and 23 and 12 miRNAs of the 191 detected miRNAs were specifically detected respectively in Col and L*er* (Additional file 2: Fig. S3c), a proportion related to the variation detected for protein-coding genes. Third, we analyzed the proportion of specific expression for the vast majority of 21/22nt and 24nt siRNAs (Fig. 3c, d) located in the different genome annotations (coding or non-coding). Altogether, the L*er* ecotype produces a larger number of 21-22nt siRNAs specifically linked to this ecotype whereas Col is more enriched for ecotype-specific 24nt siRNAs, potentially suggesting links with differential post-transcriptional and epigenetic regulations among ecotypes.

Nevertheless, the major difference in the non-coding transcriptome of the two ecotypes was linked to lncRNAs and not associated to small RNAs, even though in certain cases small RNAs may be involved in ecotype-specific regulations.

### Differential accumulation of transcripts between ecotypes along a low phosphate kinetic

The root growth arrest of Col ecotype occurred in the first hours of low phosphate sensing by the root tip [38] whereas L*er* continues its growth. To determine the effect of a short kinetics of phosphate deficiency, we examined gene expression patterns in the two ecotypes in response to this stress. Principal component analysis (PCA) showed a data dispersion that allowed a clear distinction between the effect of ecotype (first axis, Additional file 2: Fig. S4a) and the effect of the kinetics (second axis, Additional file 2: Fig. S4a). Thus, we used a multifactor analysis that takes into account the ecotype, the kinetics, their interaction and the replicate to investigate differential gene expression. Coding and non-coding genes had comparable dispersion in our experiments. Therefore we were able to use both types of gene in the same analysis. For each comparison, we confirmed the distribution of p-value as a criterion of robustness of the test [55]. After processing the differential analyses, we interpreted the results by separating the genes as “coding” or “non-coding” using the categories defined previously.

For coding genes, we observed that 3321 genes were differentially expressed between the two ecotypes on average over the 3 time points of the kinetic and 2504 were differentially expressed between at least two points of the kinetic on average over the 2 ecotypes(Fig. 4a; Additional file 1: Table S3). In our experiment, the number of differentially expressed coding genes between ecotypes or along the stress kinetics was of the same order of magnitude. However the response to phosphate starvation of only 55 genes was significantly impacted by the ecotype (“interaction” of both factors, Fig. 4a). Respectively 1566 and 1749 coding genes were up-regulated in Col and in L*er* on average over the 3 time points of the kinetic (Additional file 2: Fig. S4b). Interestingly, a clear bias of expression between ecotypes could be observed for non-coding genes (Fig. 4b). Indeed, 675 (666 + 6 + 2 +1) non-coding genes were differentially expressed between the two ecotypes on average over the 3 time points of the kinetic while only 70 (61 + 6 + 2) were differentially expressed along at least one point of the kinetic. This bias could be correlated with the number of non-coding genes specifically detected in the two ecotypes (Fig. 1a). Comparable biases were observed for both classes of non-coding genes, lincRNAs and NATs (Fig. 4c, d). Globally, 146 lincRNAs and 236 NATs were significantly up-regulated in Col compared to L*er* and 106 lincRNAs and 187 NATs in L*er* compared to Col (Additional file 2: Fig. S4c).

**Fig. 4.**
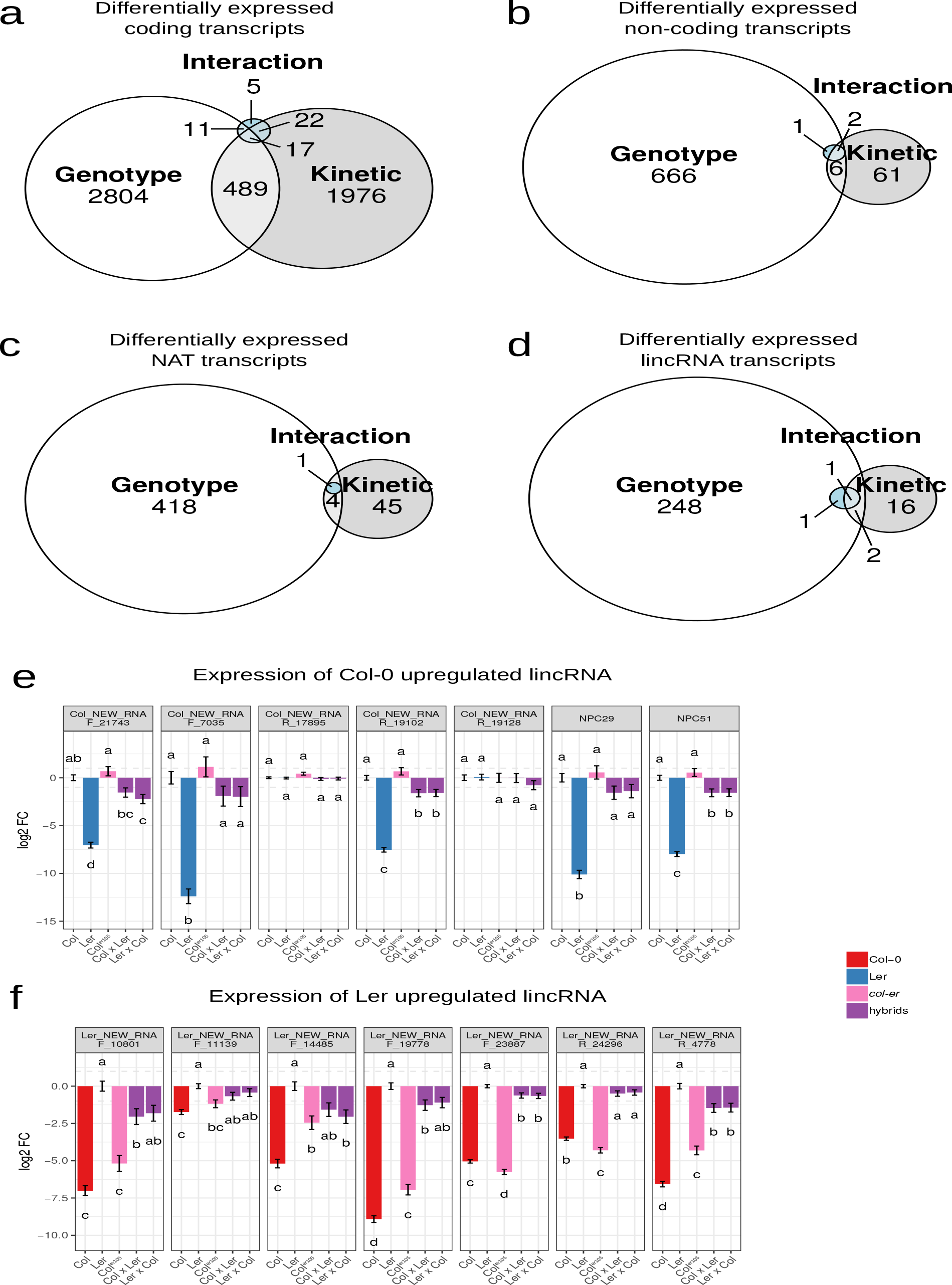
Differentially expressed gene according to ecotype and kinetic effects. Statistical analysis revealed differentially expressed genes between ecotypes and kinetics treatments for coding and non-coding genes. The differentially expressed genes can be grouped according to their significant link with genotype effect (different level between the two ecotypes), kinetic effect (differential between any couple of the points of the phosphate starvation kinetic) and their interaction (showing differential expression in response to phosphate according to the genotype). This global distribution was partitioned between coding (a) and non-coding (b) transcripts. Among non-coding genes we separated non-coding transcript antisense to another annotation (c) or intergenic ones (d). The level of expression of strongly upregulated lincRNAs was investigated by RT-qPCR in 11 days old root grown on high phosphate condition of Col, L*er*, Col^er105^ and hybrids between Col and L*er* for Col upregulated lincRNAs (e) and L*er* upregulated lincRNAs (f). Measures represent log 2 fold changes compared to Col (e) or L*er* (f). Error bars represent standard error. Results were analyzed by one-way analysis of variance (ANOVA) followed by Tukey’s post-hoc test: groups with different letters are statistically different (p ≤ 0.05) and groups with the same letters are statistically equal (p ≤ 0.05)

We used qRT-PCR in independent replicates of Col, Col^er105^ and L*er* to confirm the differential expression of 14 intergenic lncRNA genes (7 in Col and 7 in L*er*) previously identified in our statistical analysis. We were able to confirm the differential expression of 12 lncRNAs (Fig. 4 e, f). Globally Col and Col^er105^ (used for the RNA sequencing data) showed similar expression levels despite minor differences for some lncRNAs. To investigate any dominant expression effect from one ecotype, we investigated the level of expression of these lncRNAs in the F1 offsprings of Col and L*er* crosses. Among the 12 differentially expressed genes, an intermediate level of repression has always been detected (8 were statistically significant; Fig. 4e, f). This suggests independent regulation of lincRNAs between the parental genomes and discards major dominant “trans” regulatory effects of lincRNA expression between genomes.

One interesting possibility is that specific lncRNAs may be expressed in the Col and L*er* genomes in relation with known regulators of root growth responses triggered by phosphate starvation. Indeed, we could identify 2 specific antisense lncRNAs in L*er* complementary to the phosphate transporter *AT5G43370/PHT1.2* which itself is differentially expressed between ecotypes (Additional file 2: Fig. S5a). Furthermore, a Col expressed NAT RNA is complementary to *SPX4*, a critical regulator of phosphate responses, that shows reduced expression in Col compared to L*er* (Additional file 2: Fig. S5b). In other cases, we observed that two consecutive coding transcripts showing differential levels of expression among ecotypes flank an intergenic lncRNA with an ecotype-specific expression pattern (Additional file 2: Fig. S5c), suggesting that various cis-effects may be involved in these differential ecotype-linked expression patterns.

### Differential accumulations of small RNAs between ecotypes

The differential accumulations of small RNAs of 21/22nt and 24nt have been also examined in each ecotype and during the kinetics response. PCA of these sequencing data separated again clearly small RNA abundance between ecotypes but not at the level of the kinetics response (Additional file 1: Table S3; Additional file 2: Fig. S6a, b). We identified 416 coding and 211 non-coding genes that accumulated 21/22 nt siRNAs differentially between ecotypes on average over the 3 time points of kinetic with generally more siRNAs in L*er* (298 coding genes, 83 lincRNAs and 40 NATs) than in Col (118 coding genes, 49 lincRNAs and 39 NATs; Additional file 1: Table S3; Additional file 2: Fig. S6d). A greater number of genes differentially accumulated 24 nt siRNAs co-localizing with 1149 coding-genes and 429 non-coding genes between ecotypes on average over the 3 time points of kinetic, again with a greater number of differentially up-regulated siRNAs in L*er* (758 coding genes, 189 lincRNAs and 67 NATs) compared to Col (391 coding genes, 104 lincRNAs and 69 NATs; Additional file 2: Fig. S6e).

Concerning miRNA, the only difference that could be analyzed according to the PCA was also between ecotypes (Additional file 2: Fig. S6c). Indeed, among the 240 miRNAs indexed in miRBase, 38 were differentially expressed between the two ecotypes (15 and 23 for Col and L*er* respectively, Additional file 2: Fig. S6f, Additional file 1: Table S3). Interestingly, the families of miR399 and miR397 were specifically accumulated in the L*er* ecotype. These miRNAs target the PHOSPHATE 2 (PHO2) and NITROGEN LIMITATION ADAPTATION (NLA) transcripts. The PHO2 and NLA proteins are known to act together to allow the degradation of the phosphate transporter PT2 (Pht1;4) [56]. In L*er*, the higher amount of miR399 and miR397 would lead to a lower level of PHO2 and NLA and therefore a higher level of PT2 protein and could have consequently increased Pi uptake, even if the numerous post translational regulations affecting Pi transporters could limit such impact [57–60]. However, in our experiments, no difference in the accumulation of these three gene transcripts has been detected between ecotypes (as previously reported [60,61]). This suggests that the promoter activity of *PHO2* and *NLA* could enhance the transcription of these genes to compensate the increased accumulation of these miRNAs in L*er*.

### Mis-regulation of lncRNA expression affects primary root growth in Col

The different patterns of gene expression of lncRNAs between ecotypes may be relevant for the generation of novel regulatory patterns linked to root growth responses. As the interactions of lncRNAs with different ribonucleoproteins may lead to changes in gene expression potentially linked to the differential growth responses between Col and L*er*; we selected 5 lncRNA genes: *NPC15, NPC34, NPC43, NPC48* and *NPC72* showing differential expression among ecotypes in order to study the impact of their expression on Col primary root growth. In our RNA-Seq data, three of these lncRNA genes were more expressed in Col (*NPC15, NPC43* and *NPC72*) and two in L*er* (*NPC34* and *NPC48*) (Fig. 5a). To evaluate the potential dominant expression patterns of these lincRNA genes, we investigated their expression in corresponding two F1 of reciprocal crosses Col x L*er*. Expression analysis by qRT-PCR confirmed the previous RNA-Seq expression levels for *NPC15, NPC34* and *NPC72* genes (Fig. 5b) whereas for NPC48, the differential expression was only detected in the Col^er105^ mutant (the line used for the RNA-Seq experiment), suggesting that the erecta mutation directly affects the expression of this gene. No differential expression could be detected for the NPC43 gene. This latter result might be due to the accumulation of antisense transcripts (*NPC504*) that could generate siRNAs against NPC43 transcripts. One explanation for the differential expression of lincRNA genes in these two ecotypes could be linked to genetic changes at the transcript locus. Hence, we examined the DNA sequences at these specific loci (Additional file 2: Fig. S7; profiting from the well characterized Col and L*er* genomes). No significant modifications (except few SNPs) were detected between Col and L*er* for the *NPC34, NPC43* and *NPC48* loci. By contrast, the *NPC15* locus contains an insertion of 2417 nt in L*er* v8 and *NPC72* is completely missing in L*er* v7 and v8 genomes (Additional file 2: Fig. S7a-e). Therefore, genome modifications could explain the specific expression pattern of *NPC72* and *NPC15* genes in L*er*.

**Fig. 5.**
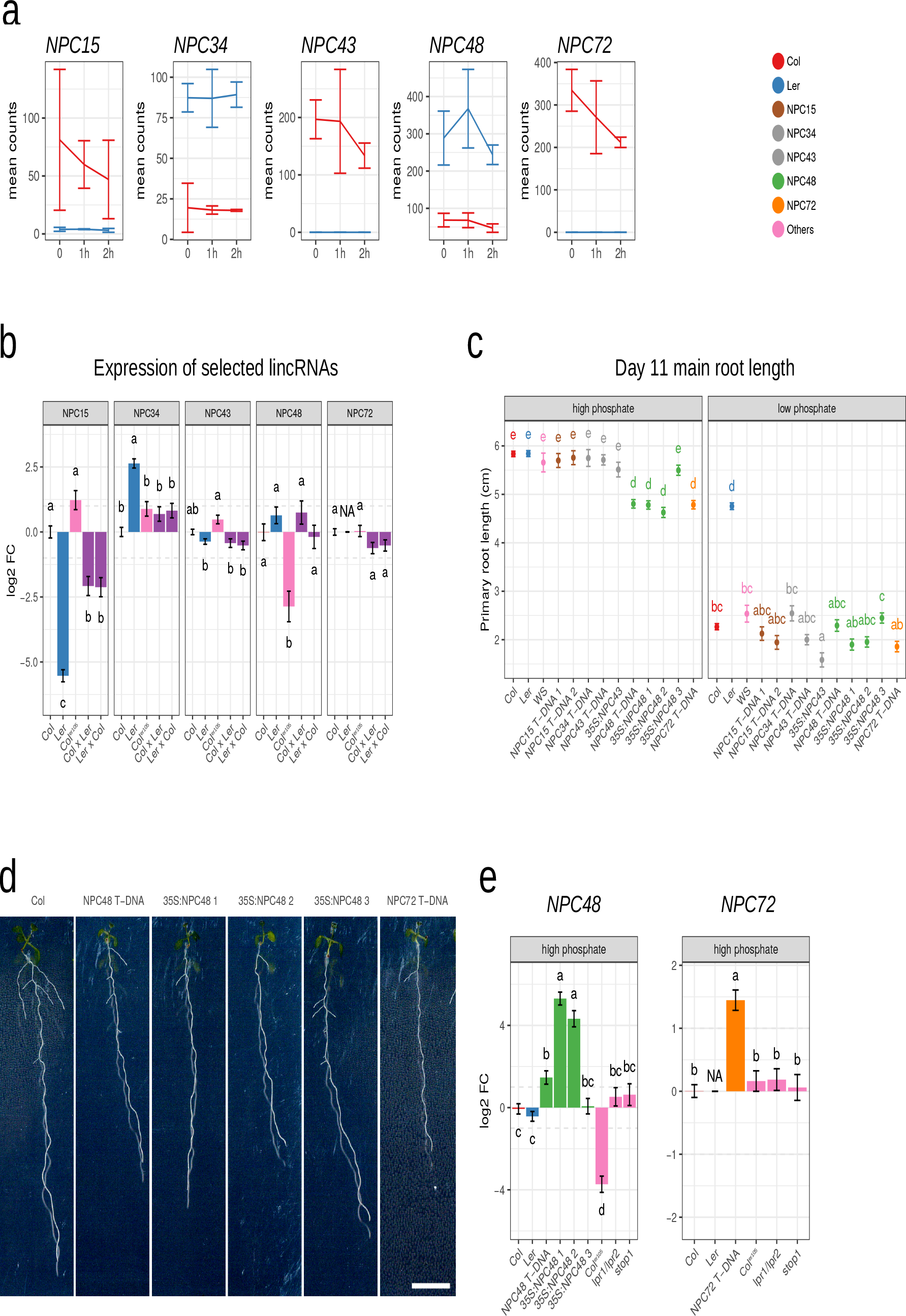
Overexpression of the lincRNAs *NPC48* and *NPC72* affects primary root growth. (a) Expression profile of selected intergenic lincRNAs in Col and L*er* early phosphate starvation kinetics (RNAseq data, average expression and standard deviation). Five selected lincRNAs showed differential expression at each time point between Col and L*er*. (b) Level of expression of selected intergenic lincRNAs in 11 days old root grown on in high phosphate condition in Col, L*er*, *Col^er105^* and hybrids between Col and L*er*. Measures represent log 2 fold changes compared to Col measured by RT-qPCR. (c) Mean primary root length according to genotype and phosphate condition at the age of 11 days after sowing. (d) Expression level of *NPC48* and NPC72 in 11 days old root grown on high phosphate condition of lines deregulated in *NPC48* or *NPC72* and mutants affected in phosphate related root arrest. Measures represent log 2 fold changes compared to Col. (b) - (d) Measure represent corrected means of the FC (b) and (d) or of primary root growth (c) computed according to generalized linear model fitted on several experiments (see the “Methods” section) and taking into account the following factors: genotypes, growth condition, interaction between genotype and growth condition. Error bars represent standard error. Results were analyzed by one-way analysis of variance (ANOVA) in (b) and (d) or two way ANOVA in (c) followed by Tukey’s post-hoc test: groups with different letters are statistically different (p ≤ 0.05) and groups with the same letters are statistically equal (p ≤ 0.05)

To support the biological relevance of lincRNAs differentially regulated between ecotypes at phenotypic level, we used T-DNA insertion and over-expressing (OE) lines to monitor the effect of lincRNA genes on Col root growth in control and low phosphate conditions. In control conditions, only *NPC48* and *NPC72* overexpression lead to a significant root growth reduction when compared to Col (Fig. 5c). Indeed, T-DNA insertions have been mapped to the 5’ region of the *NPC48* and *NPC72* loci respectively for *npc48* T-DNA and *npc72* T-DNA and lead to an overexpression of these lincRNAs in the respective lines (Fig. 5d). For npc48 T-DNA, it strongly supported the phenotype observed with the 35S::NPC48 lines. In low phosphate conditions, known to inhibit root growth in Col but not in L*er*, minor differences in root length were observed. However, the OE of *NPC43* gene clearly increased the inhibition of root growth (Fig. 5c) when compared to Col plants. A *priori*, the ratio of root growth in control and low phosphate conditions should highlight potential phenotypic differences in phosphate sensitivity of transgenic lines. This ratio is significantly increased for **NPC48** and *NPC72* lines compared to Col, partly mimicking a L*er*-type response. On the other hand, this ratio is significantly reduced for NPC43-and NPC15-deregulated lines (Fig. 5c).

*NPC48* and *NPC72* de-regulated lines present a significant decrease of root length in control, but not in low phosphate conditions. The 500 μM of Pi used in our control conditions could be perceived as a low Pi concentration in these lines. Hence, we asked whether these phenotypes could be linked to a root growth arrest due to oversensitive perception of phosphate starvation under control conditions or an alteration of Pi systemic sensing (which would affect Pi uptake). This does not seem the case as i) known phosphate-starvation markers were not induced in root in control conditions and ii) these markers were induced in these lines at the same level as in Col (Additional file 2: Fig. S8a, b). Then we investigated the local Pi signaling response which control primary root growth. As the genes *LOW PHOSPHATE ROOT 1* and *2* (LPRs) and the transcription factor STOP1 are known to be involved in primary root growth arrest under low Pi [38] we analyzed the expression of *NPC48* and *NPC72* genes in *lpr1*/*lpr2* and stop1 mutant lines. Fig. 5e shows that we cannot observe any significant variation of these lncRNA expression patterns in these mutant lines. Reciprocally, no significant expression variation was detected of the LPR1/LPR2 pathway in *NPC48* or *NPC72* lines (*LPR1, LPR2, STOP1, ALMT1* and *MATE* genes; Additional file 2: Fig. S8c-g).

Altogether, the identification of ecotype-related lncRNAs allowed us to characterize new regulators of primary root growth acting through a distinct pathway from that involving *LPR1/LPR2* and *STOP1*.

## Discussion

Until recently, transcriptome studies were mainly focused on protein coding gene transcripts and discarded lncRNAs. Variation in the nucleotide sequences or expression patterns of the non-coding genome can have less pleiotropic effects than changes in the protein sequence of critical regulators. Therefore, in addition to promoters, introns and transposons, regions encompassing non-coding RNAs emerge as actors of plant adaptation to environmental constraints. It is now clear that certain lncRNAs play important roles in development and response to environmental conditions [1], even if the majority of lncRNAs awaits a functional characterization. In addition, lncRNAs exhibit a high cell specificity which could explain specific functions in a particular cell-type in plants and animals. In this study, by using strand specific RNA analysis on root tips, we succeeded in identifying thousands of different forms of lncRNAs: lincRNAs, NATs or antisense RNAs expressed from two Arabidopsis accessions, Col and L*er*. As previously shown in numerous studies, in comparison with mRNAs, lncRNAs observed here were expressed at low levels [62,63]. We opted to focus on root growth as it is a complex trait resulting of the expression of a large number of loci spread across the plant genome and is highly susceptible to the soil environment [64]. Interestingly, in our study, we observed many lncRNA genes differentially or specifically expressed in Col or L*er*, in contrast to protein-coding genes for which the expression were more stable in the two ecotypes. We think reasonable to assume that certain of these lncRNAs could contribute to Col and L*er* root growth specificities. Indeed, in animals, certain complex traits are mainly driven by non-coding variants [65,66]. In chicken, domestication traits governing body morphology and behavior are under selection and correspond to lincRNA genes [67]. Moreover, recently, the human LincSNP 2.0 database, which contains 809451 unique disease associated SNPs, 11,6 million of linkage disequilibrium SNPs and 244 545 lncRNAs, identified approximately 45% of disease–associated human SNPs that mapped to non-coding regions of the genome [68]. Some lncRNAs containing SNPs have been recently associated to cardiometabolic traits [69]. Similarly, in plants, the comparison of SNPs observed in fruit transcripts of two tomato cultivars also corresponded to non-coding genomic regions or lncRNA genes [70]. The SNPs could act directly at the level of lncRNA expression or affect the expression of lncRNA-neighboring genes [24,71]. LncRNAs are thus elements to be considered in genetic association studies.

Root apexes play an important role in sensing external stimuli. We examined the gene expression profile soon after the stress application (1 h and 2 h) in two ecotypes that present different root growth phenotypes in response to phosphate starvation. The number of coding genes differentially expressed during the phosphate kinetic and between the two ecotypes was of the same order of magnitude. A clear bias of specific expression of lncRNA genes was identified between the two ecotypes. This ecotype bias was also observed for the accumulation of siRNA. The analysis of pan-genome (restricted to coding genes) using genomic and RNAseq of 19 Arabidopsis ecotypes showed that at least 70 accessory genes could be identified in each ecotype [25]. In response to stress, it is known that accessory genes can explain, at least, part of the phenotypic difference of behavior observed among ecotypes [44]. In our experiments, the expression polymorphism (differential gene expressions observed between Col and L*er*) corresponded mainly to regions where SNPs accumulate or neighboring DNA rearrangements (e.g. transposon insertions) were detected. Very few ecotype-specific lncRNAs coincided with the absence of specific DNA sequences in the particular ecotype. Hence, we propose that the lncRNA difference of expression of a relatively similar DNA molecule would result in shifts in transcription or stability of lncRNAs that could be connected to SNP or InDels polymorphisms in promoters and/or lncRNA gene sequences in the two ecotypes. As lncRNAs can repress or activate the transcription of other genes the expression polymorphisms observed between the two ecotypes could also result in a cascade of cis-local or trans-distal action on target genes [72,73]. It is noteworthy that the majority of ecotype-specific lncRNAs identified did not co-localize with siRNAs and thus could not reflect only gene silencing differences between ecotypes (either transcriptional or post-transcriptional processes, [53,54]). This points to the lncRNA itself or its transcription *per se* to be linked to the quantitative regulation of target gene expression [73].

We were able to confirm, by qRT-PCR, the expression of 12 lincRNAs among the 14 chosen for validation, supporting the expression polymorphisms identified. Allele-specific expression is known to affect specific diseases in human and productivity in plant and animal agriculture [74,75]. Moreover, in Arabidopsis, heterosis has been reported for different traits such as flowering time [76] leaf area and plant biomass [77] but also for phosphate acquisition [78]. As lncRNAs are able to modify chromatin and thus alter gene expression, we added, in our expression analysis, F1 resulting from reciprocal crosses between Col and L*er*. In the F1 offspring, the 15 confirmed differentially expressed lncRNA genes chosen for validation exhibited globally an additive expression pattern when compared to their parents. This is consistent with results obtained in maize F1 hybrids, where additivity are frequently observed for lncRNAs [79] and 78% of coding genes [80]. In the case of non-additive expression patterns in the heterozygote, a *trans*-regulation could affect the two alleles and products a specific accumulation of lncRNA transcripts.

The general adaptation of root architecture in response to low Pi comprises an arrest of PR growth and the activation of LR development after the perception of Pi limitation. In Arabidopsis, studies concerning plant phosphate homeostasis, during Pi deficiency, characterized the IPS1/AT4 lncRNA controlling the distribution of Pi from root to shoot. It acts as a target mimic for miR399, which regulates *PHO2* mRNAs [81,82]. Moreover, Yuan et al [62] identified lncRNAs differentially expressed in root and shoot of plants grown in the presence or absence of Pi for 10 days. The authors suggested that a co-expression between lncRNAs and adjacent coding genes may be linked to a cis-regulation by lncRNAs of target genes involved in Pi starvation processes. Interestingly, in fission yeast *S. pombe*, two of the three genes of the phosphate regulon are repressed, in Pi rich medium, by the transcription of lncRNA genes [83,84] that are present in the 5’ region (cis-regulation). The molecular mechanisms that govern root growth modification by Pi have been mostly elucidated in Col plants. For the local impact of Pi (restricted to root architecture), Pi deficiency is sensed by the root tips and a primary root growth inhibition is induced by both the reduction of cell elongation and the progressive arrest of meristem division, notably linked to the presence of iron in the medium. The pathway that governs this meristem arrest involves the STOP1 transcription factor that controls the expression of the transporter ALMT1 involved in malate secretion [38]. In the apoplasm, malate leads in turn to the production, by LPR1/LPR2, of ROS that induces plasmodesmata (PD) closure by callose deposition [40]. The interruption of trafficking through PD progressively blocks meristem division. In parallel, in low Pi, it has been shown recently that the small peptide CLE14, which acts after LPR1/LPR2 module, is able to trigger the differentiation of root meristem in absence of callose deposition [37]. We selected several ecotype-specific lncRNAs for functional root growth analysis and misregulation of two lncRNAs affected primary root growth. Expression analysis in response to Pi deficiency in our mutant lines *NPC48* and *NPC72*, allowed us to suggest the existence of a potential new pathway that do not overlap with the known LPR1/LPR2 and STOP1 pathways. In Arabidopsis, by using grafts between ecotypes presenting a high frequency of SNPs, Thieme et al [85] have shown that about 2000 mRNAs, among 9300 containing SNPs, could move in plants that were subjected to Pi deficiency for two weeks. These mRNAs were transported from root-to-shoot or shoot-to-root. The authors suggested that these mobile mRNAs might function widely as specific signaling molecules coordinating growth, cell differentiation and stress adaptation of distant body parts. As the lncRNAs described here have 3’ polyA tails and are probably 5’ capped, it is tempting to assume that at least some of them can be transported through the xylem and/or the phloem and may contribute to systemic signaling responses.

Globally, the in depth exploration of the non-coding transcriptome of two ecotypes identified thousands of new lncRNAs with ecotype-specific expression. Statistical analysis among ecotypes identified several co-regulated events between coding and non-coding genes (including small RNAs) potentially linked to the evolution of different regulatory mechanisms among ecotypes grown in diverse soil environments. De-regulation of two ecotype-related lncRNAs revealed a new pathway involved in the regulation of primary root growth.

## Methods

### Plant growth

Seeds were surface-sterilized seeds were sown on a horizontal line in plates that were vertically disposed in a growing chamber (16 h photoperiod; intensity 90 µE; 21 °C). The growth medium contained 0.15 mM MgSO_4_, 2.1 mM NH_4_NO_3_, 1.9 mM KNO_3_, 0.34 mM CaCl_2_, 0.5 μM KI, 10 μM FeCl_2_, 10 μM H_3_BO_3_, 10 μM MnSO_4_, 3 μM ZnSO_4_, 0.1 μM CuSO_4_, 0.1 μM CoCl_2_, 0.1 μM Na_2_MoO_4_, 0.5 g.L^-1^ sucrose. The agar (0.8 g.L^-1^) for plates was from Sigma-Aldrich (A1296 #BCBL6182V). The -Pi and +Pi agar medium contained 10 and 500 μM Pi, respectively; the media were buffered at pH 5.6-5.8 with 3.4 mM 2-(N-morpholino)ethane sulfonic acid.

### Arabidopsis lines

The stop1 (SALK_114108, NASC reference N666684), lpr1;lpr2 [36], npc15 T-DNA 1 (SALK_027817, NASC reference N527817) npc15 T-DNA 2 (SALK_090867, NASC reference N590867), npc43 T-DNA (SALK_007967, NASC reference N507967), *npc48* T-DNA (SAIL_1165_H01, NASC reference N843057) and npc72 T-DNA (SAIL_571_C12, NASC reference N824316) are in the Col-0 (Col) background. Surexpressor lines 35S:NPC43, 35S:NPC48 1, 35S:NPC48 2, 35S:NPC48 3 were retrieved from [49] and are in Col-0 background. npc34 T-DNA (FLAG_223D08 or FLAG_228A07) is in WS background. Col^er105^ is in a Columbia background (Col-0) with the null allele erecta-105 [86]

### Libraries construction and sequencing

Three biological replicates of Col^er105^ and L*er* were sown vertically on 1 cm high bands of nylon membrane (Nitex 100 μm). After one week on +Pi agar medium, while the roots were out of the membrane, the membranes were transferred on -Pi agar medium. Plants were then sampled at time point 0, 1 h and 2 h after transfer. Each biological replicate is a pool of more than 100 root apexes cut at 0.5cm from root extremity. Total RNA were extracted using RNeasy micro kit (Qiagen 74004, n°lot 136257409), tissues protocol. One microgram of total RNA from root tips of each sample was used for mRNA library preparation using the Illumina TruSeq Stranded mRNA library preparation kit according to the manufacturer instruction. Libraries were sequenced on HiSeq™ 2000 Sequencing System (Illumina) using 100 nt paired-end reads. Samples were multiplexed by 6 and sequenced on 4 sequencing lines. All reads were quality trimmed using Trimmomatic and remaining ribosomal sequences were removed using sortMeRNA [87].

For small RNAs libraries, root apexes grown and collected in the same condition as for the RNA-Seq were used. Small RNAs were extracted using mirVana™ miRNA Isolation Kit (Ambion AM 1560, part IV/A. Isolation of Small RNAs from Total RNA Samples). Small RNA libraries were construct using Ion Total RNA-Seq Kit v2 (Ion Torrent, Life Technologies) according to manufacturer instruction. Libraries were then sequenced using IonProton and the adapters removed.

### New transcript identification

According to their ecotype of origin, mRNA cleaned reads were aligned on TAIR10 [43] or L*er* v7 [44] genome using tophat2 (version 2.0.13, [88]) with the following arguments: --max-multihits 1 --num-threads 8 -i 20 --min-segment-intron 20 --min-coverage-intron 20 --library-type fr-firststrand -- microexon-search -I 1000 --max-segment-intron 1000 --max-coverage-intron 1000 --b2-very-sensitive. Independently of ecotype, new transcripts were predicted using GFFprof included in RNAprof [89]

The transcripts predicted on each genome, TAIR10 and L*er* v7, were positioned on the other genome, respectively L*er* v7 and TAIR10, and on L*er* v8 [45] using blastn (from BLAST suite 2.2.29+) using a maximum e-value of 10^-4^. For each transcript, the different blast hits fragments were fused together if the distance between two fragments was less than 5000 nucleotides and placed on the same strand of the chromosome. Only hits with at least 90 % of sequence identity and where the length was conserved (at least 90 % and less than 110 % of length outside of insertion) were kept. For each transcript only the best hit was conserved according first to the conservation of the sequence length and then identity. In case of hits of the same strength, a higher priority was given when the chromosome and then the strand were conserved. Each transcript was therefore placed on each of the three genomes.

For L*er*-predicted transcripts only those positioned on L*er* v8 were kept. On each genome independently, transcripts coming from the same ecotype (GFFprof prediction) or the other one (blastn positioning) were fused using cuffmerge (version 1.0.0) with default parameters. Only transcripts longer than 200nt in either ecotype were kept for further processing.

Based on the position of the transcripts on the TAIR10 genome, new transcripts were annotated according to already known transcripts in the following databases: Araport 11 [46], RepTas [8], CANTATAdb [47], miRBase v21 [48] lncRNAs predicted from Ben Amor et al. [49] and root predicted lncRNAs from Li et al. [50]. GffCompare (version 0.10.4, https://ccb.jhu.edu/software/stringtie/gffcompare.shtml) was used for the comparison. In case of overlap with a known transcript (=, c, k, j, e, o codes of GffCompare), the closest transcript was used to determine the identification and the coding potential of the transcript. For previously non-discovered transcripts we used the COME software [90] to predict their coding potential.

### Library saturation analysis

The saturation of libraries was computed using the `RPKM_saturation.py` script included in RSeQC v3.0.0 [91,92]. Raw counts were quantified at gene level from resampled bam for each additional 2% of the reads from 2% to 100% of the reads (`-l 2 –u 100 –s 2`). The saturation was computed 100 times. For each library and each sampling the number of detected gene (known or new and coding or non-coding) was defined as the number of genes having at least one read.

### Small RNA analysis

The cleaned small RNA reads were aligned on TAIR10 or L*er* v7 genome using ShortStack (version 3.8.5, [93]) without mismatch (`--mismatches 0`), keeping all primary multi-mapping (`--bowtie_m all`) and correcting for multi-mapped reads according to the uniquely mapped reads (`--mmap u`).

For each annotation in Araport11 (mRNA coding and non-coding and TE) and each new annotation predicted in this study according to mRNA sequencing the accumulation of small RNA was analysed using ShortStack with default parameters. The counts for 21nt and 22nt were summed for each sample. Using these new counts, the DicerCall, defined as the size of the majority of the reads of a cluster, was recomputed. The other description and counting are according to ShortStack prediction.

### Expression analysis

For each annotation, coding and non-coding, mRNA reads were counted with htseq-count [94] using strand specific and intersection strict mode (`-- stranded=reverse -t gene --mode=intersection-strict`). These counts were used for differential gene expression analysis with DEseq2 (v1.16.1[95]) using a linear model and as factors the ecotype (two levels), the kinetic time (3 levels), the interaction between the two and the replicate (3 levels). Low counts were discarded using the default DESeq2 threshold and raw p-values were adjusted with the Bonferroni method. Differentially expressed genes were defined as having an adjusted p-value lower than 0.01.

Differential siRNA accumulation was computed using DESeq2 with a model taking into account only the genotype (two levels) as factor, and using the counts of ShortStack. Differential accumulation was computed independently for the 21/22nt on one side and the 24nt on the other side and limited to coding and non-coding genes.

Bonferroni correction of the p-value was used and differential siRNAs were defined as having an adjusted p-value inferior to 0.01.

### Measurement of the primary root length

Pictures of the plates were taken with a flat scanner and root lengths measured using RootNav software [96]. Each measure corresponds to a different plant.

### Quantitative RT–PCR

Total RNA was extracted from whole roots using the Quick-RNA MiniPrep™ kit (Zymo Research, USA) and treated with the included DNAse treatment according to the manufacturer’s instructions. Reverse transcription was performed on 500 ng total RNA using the Maxima Reverse Transcriptase (Thermo Scientific™). Quantitative PCR (qRT– PCR) was performed on a 480 LightCycler thermocycler (Roche) using the manufacturer’s instructions with Light cycler 480sybr green I master (Roche) and with primers listed in Additional file 1: Table S4. We used PP2A subunit PD (AT1G13320) as a reference gene for normalization.

### Statistics and reproducibility of experiments

Statistical analyses were performed using R (v3.4.2 [97]) with the help of the tidyverse (v1.2.1 [98]) and `emmeans` packages [99]. For each measure (root length, qPCR expression level, experimental repetition), the least-squares means were computed taking into account all the factors (genotype and condition) in a linear model. This allows correcting for inter-repetition variation. Data are presented as least-squares means±SEM.

The results for statistical significance tests are included in the legend of each figure. ‘n’ values represent the number of independent samples in a repetition, i.e. the number of roots or pools of root per condition. The number of independent experiments is denoted as “repetition”. For each analysis, the detail number of repetition and number of sample per repetition are available in Additional file 1: Table S5 and Table S6.

## Declarations

### Availability of data and material

Sequence files generated during this study have been deposited into the NCBI GEO database [100] under the accessions GSE128250[101] and GSE128256[102].

### Ethics approval and consent to participate

Not applicable.

### Consent for publication

Not applicable.

### Competing interests

The authors declare that they have no competing interests.

## Funding

This work was supported by grants from Agence Nationale pour la Recherche (ANR) RNAdapt (grant no. ANR-12-ADAP-0019), SPLISIL (grant no. ANR-16-CE12-0032) and grants of The King Abdulla University of Science and Technology (KAUST) International Program OCRF-2014-CRG4 and ‘Laboratoire d’Excellence (LABEX)’ Saclay Plant Sciences (SPS; ANR-10-LABX-40).

## Authors’ contributions

TB, CB, TR, LS performed gene expression analysis; TB, MG, DG, ED and MLM performed statistical analysis and bioinformatics; CB, CS and AC were involved in sample preparation and processing; TB, CH, TD, LN and MC directed experimental work; TB, CH, TD and MC designed experiments and wrote the manuscript. All authors provided comments to the manuscript and approved it.

### Acknowledgement

We thank Ambre Miassod and Janina Lüders (IPS2, CNRS, INRA, Université Paris-Sud, Université Evry, Université Paris-Saclay, France) for technical assistance with gene expression experiments.

## Additional files

### Additional file 1

supplemental tables.

- Table S1 - Mapping efficiency for each sequence sample
- Table S2 - Genomic information of new transcripts compared to TAIR10
- Table S3 - Differential gene expression analysis
- For each comparison, the list of genes differentially expressed
- Table S4 - Sequence of primers used in this study.
- Table S5 - Number of samples used for each genotype and condition in the qPCR experiments
- Table S6 - Number of samples used for each genotype and condition in root length measurements

### Additional file 2

supplemental figures:

- Fig. S1 - Characteristic of identified transcripts
- Fig. S2 - Expression level and detection of coding and non-coding genes
- Fig. S3 - Ecotype specific classification of lncRNAs as siRNA precursors
- Fig. S4 - The ecotype effect on gene expression
- Fig. S5 - Genome organization and correlation of expression at selected loci
- Fig. S6 - The ecotype effect on siRNA accumulation
- Fig. S7 - Genome homology at selected ncRNAs loci.
- Fig. S8 - Deregulation of NPC48 and NPC72 do not change the expression of phosphate starvation related genes.

### Additional file 3

new transcript localization on Col and Ler v7 genome as GFF files.

**Fig. S1 - Characteristic of identified transcripts**

(a) Flowchart of identification of lncRNAs responsive to Pi starvation in two Arabidopsis ecotypes. Plants of the two ecotypes were grown on control condition for seven day before transfer on low phosphate condition. Root tip were then sampled at time point 0 h, 1 h and 2 h after transfer. After RNA extraction, PolyA transcripts were sequenced. The retrieve reads were then mapped independently for the two ecotypes on their respective genomes (TAIR10 for Col and L*er* v7 for L*er*). Based on this mapping we predicted new transcriptional units on each genome compared to TAIR10 annotation available on both genomes. We then aligned the resulting transcriptional on the opposite genome to compute a homology and fused the overlapping transcripts. Only the transcript with a length superior to 200 nt were kept for further analysis. Each transcript was then categorized in one of the 4 following classes: coding, structural RNA (rRNA, tRNA, snRNA, snoRNA,…), transposable element and non-coding RNA. The classification was estimated based first the overlap with already annotated transcripts (by order of importance in Araport11, RepTAS database, CANTATA database, BenAmor et al. 2009, Li et al. 2016 and miRBase v21). For the transcripts that were not found in any of these databases, their coding potential was predicted using COME. Repartition of the different detected genes in the different Arabidopsis databases for genes predicted as coding (b) or as non-coding (c).

**Fig. S2 Expression level and detection of coding and non-coding genes**

Distribution of coding genes, non-coding genes, structural RNA and transposable element according to their level of expression in Col (a) or L*er* (b). (c) Number of newly detected genes per percent of additional sequencing reads of the library. Line correspond to the median of 100 bootstraps, grey shadow correspond to the standard deviation.

**Fig. S3 - Ecotype specific classification of lncRNAs as siRNA precursors**

ShortStack classification of all non-coding genes as siRNA precursor expressed at more than 1 RPM of major siRNA in Col (a) or L*er* (b). (c) Detection specificity of non-coding RNA as phased 21 nt and 22 nt siRNAs precursors at the level of 1 RPM in each ecotype. (d) Detection specificity of miRBase miRNAs at the level of 1 RPM in each ecotype. The majority of the miRNAs are detected in both ecotypes.

**Fig. S4 - The ecotype effect on gene expression**

(a) PCA analysis showing the effect of ecotype and phosphate kinetic on the variance of gene expression. For the 3321 coding genes (b) and 675 non-coding genes (c) that are ecotype differentially expressed, the number of genes that accumulated more in Col or L*er*.

**Fig. S5 - Genome organization and correlation of expression at selected loci**

(a) Increased expression of *PHT1;2* in L*er* correlated with the specific expression of the two NATs *Ler_NEW_R_34181* and *Ler_NEW_R_34180* (b) Decreased expression of *SPX4* gene in Col correlated to the expression of the NAT *Col_NEW_RNA_R_29088* (c) Differential expression of *NIP3;1* and *AT1G1910* between Col and L*er* correlated with the expression of lincRNA *At1NC041650*.

**Fig. S6 - The ecotype effect on siRNA accumulation**

PCA analysis showing the effect of genotype and phosphate kinetic on the variance between samples for 21/22nt (a), 24nt (b) and miRNAs (c). The samples can be well separated according to genotype (Col or L*er*) but not according to the phosphate kinetic point. For ecotype differentially accumulated siRNAs, the number of differentially accumulated siRNAs in either ecotype for 21/22nt siRNA precursors (d), 24nt siRNA precursors (e) and miRNAs (f).

**Fig. S7 - Genome homology at selected ncRNAs loci.**

Genome alignment around *NPC15* (a), *NPC34* (b), *NPC43* (c), *NPC48* (d) and *NPC72* (e) between the reference sequence of Arabidopsis Col and two sequences from L*er* (L*er* v7 and L*er* v8). The identity line is color coded according to conservation: green full conservation, yellow mismatches. For each sequence, dash represent missing sequences, plain grey identity sequence and plain black specific insertion or mismatches.

**Fig. S8 - Deregulation of *NPC48* and NPC72 do not change the expression of phosphate starvation related genes.**

(a) - (g) Level of expression in 11 days old root grown on in high phosphate condition or low phosphate condition of genes involved phosphate sensing, *IPS1* (a) and *SPX3* (b), or in the response to phosphate related growth arrest, *LPR1* (c), *LPR2* (d), *STOP1* (e), *AMLT1* (f) and *MATE* (g). Measures represent log 2 fold changes compared to Col measured by RT-qPCR. Measure represent corrected means of the FC computed according to generalized linear model fitted on several experiments (see the “Methods” section) and taking into account the following factors: genotypes, growth condition, interaction between genotype and growth condition. Error bars represent standard error. Results were analyzed by two-way analysis of variance (ANOVA) followed by Tukey’s post-hoc test: groups with different letters are statistically different (p ≤ 0.05) and groups with the same letters are statistically equal (p ≤ 0.05)

**Table S1 - Mapping efficiency for each sequence sample**

**Table S2 - Genomic information of new transcripts compared to TAIR10**

**Table S3 - Differential gene expression analysis**

For each comparison, the list of genes differentially expressed

**Table S4 - Sequence of primers used in this study.**

**Table S5 - Number of samples used for each genotype and condition in the qPCR experiments**

’n’ values represent the number of independent samples in a repetition, i.e. the number of pools of root per genotype and condition. The number of independent experiments is denoted as “repetition”.

**Table S6 - Number of samples used for each genotype and condition in root length measurements**

’n’ values represent the number of independent samples in a repetition, i.e. the number of root per genotype and condition. The number of independent experiments is denoted as “repetition”.

