## Supplementary figures and images for "Two ecotype-related long non-coding RNAs in the environmental control of root growth"

### FigS1.pdf

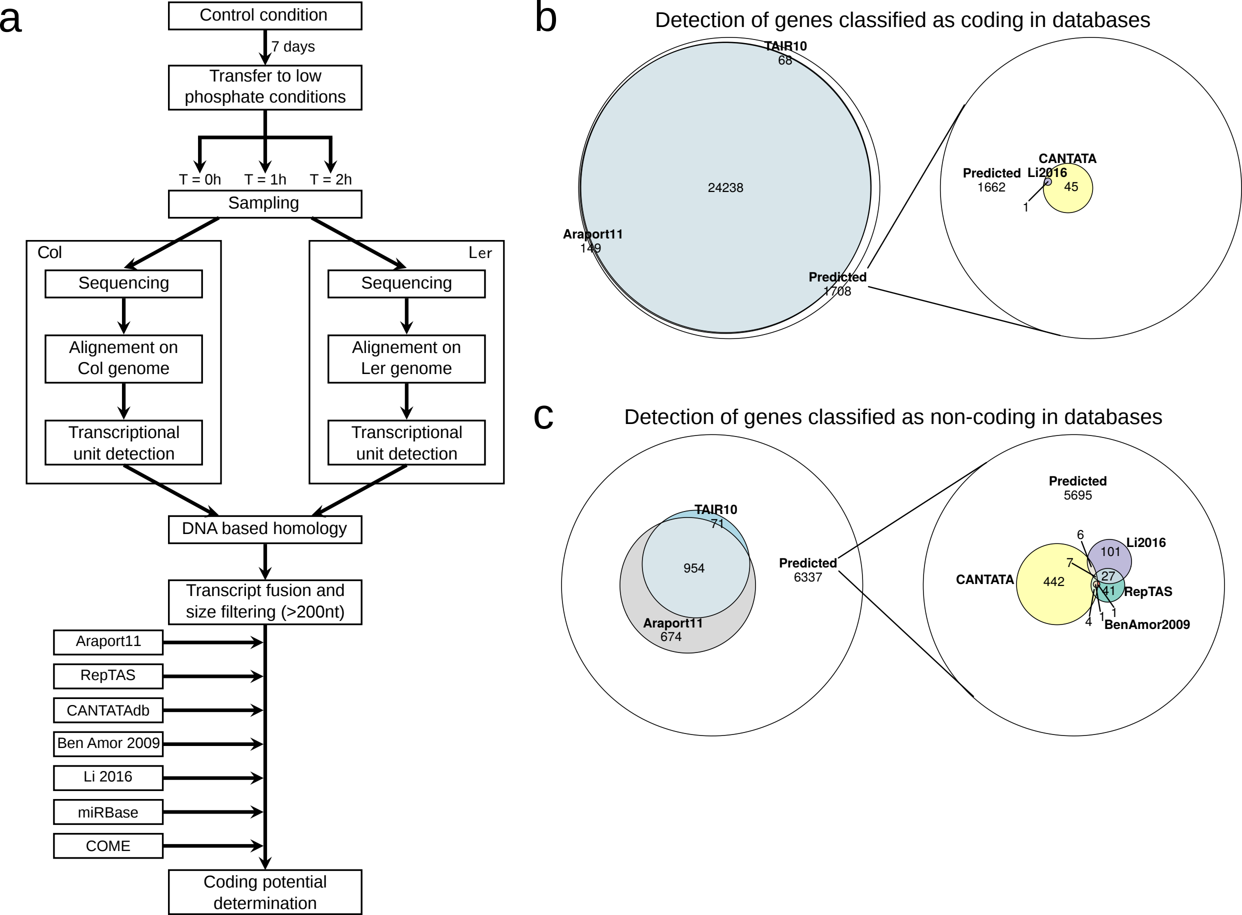

### FigS2.pdf

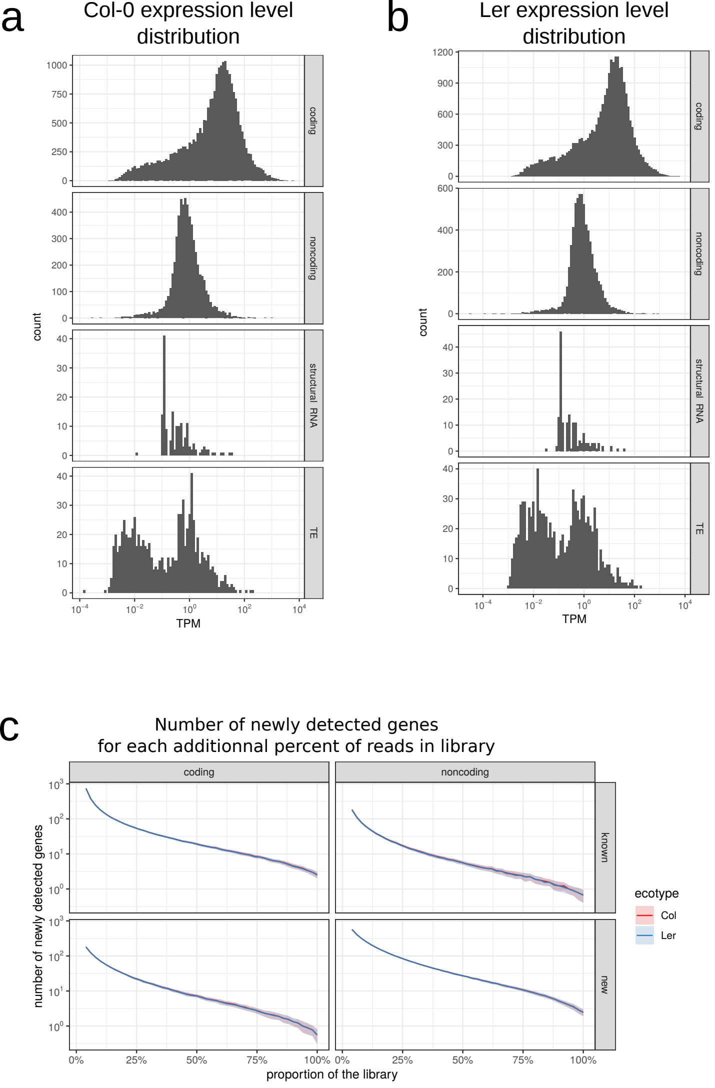

### FigS3.pdf

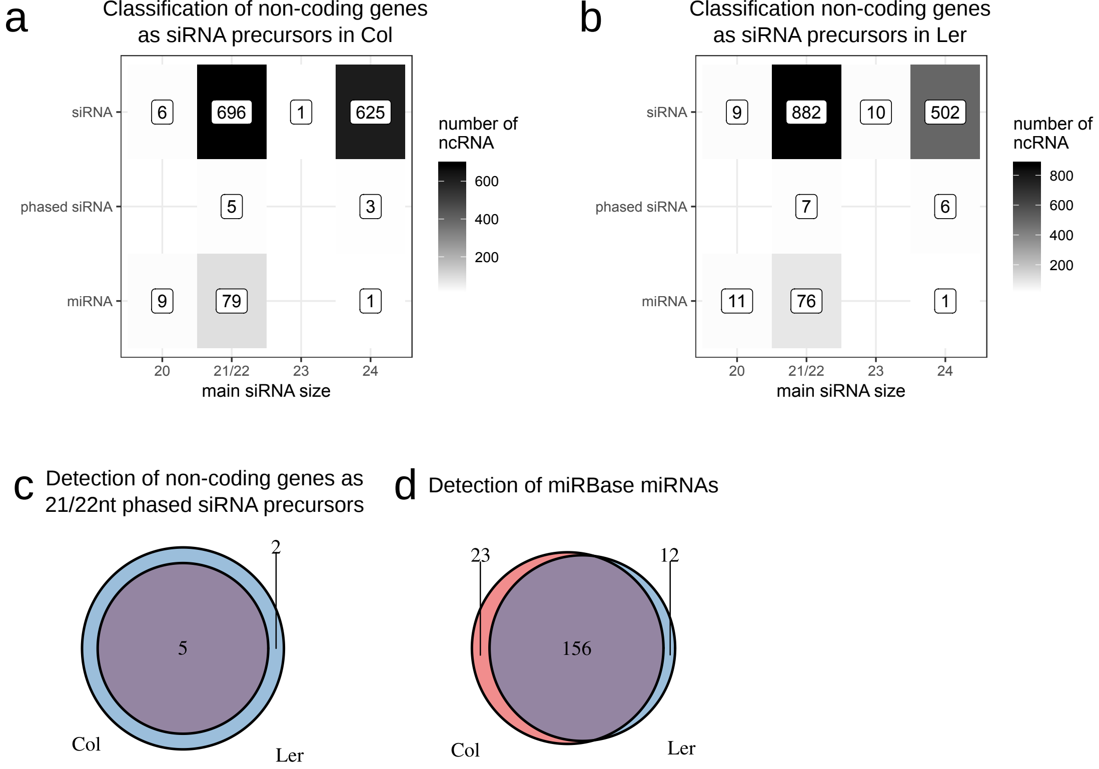

### FigS4.pdf

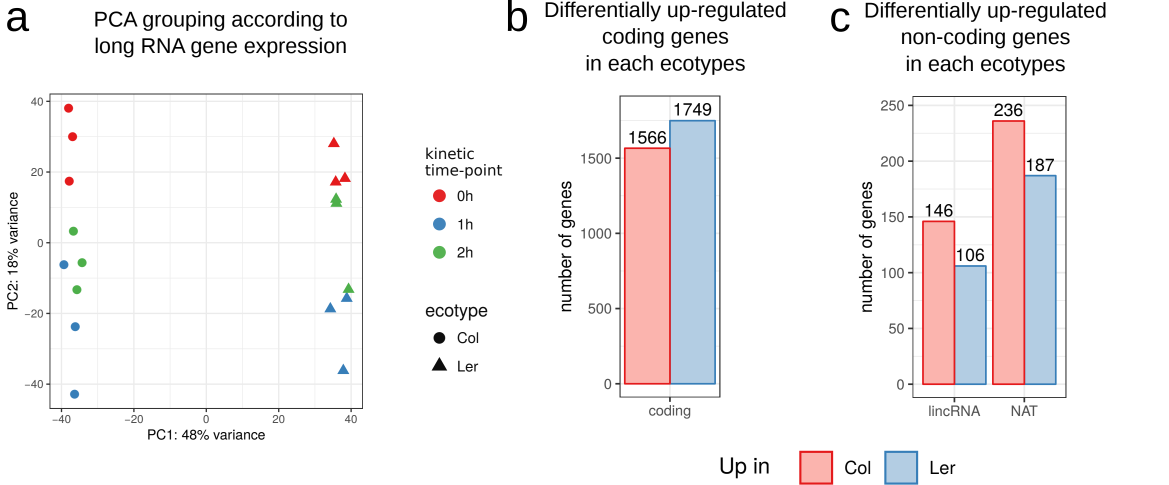

### FigS5.pdf

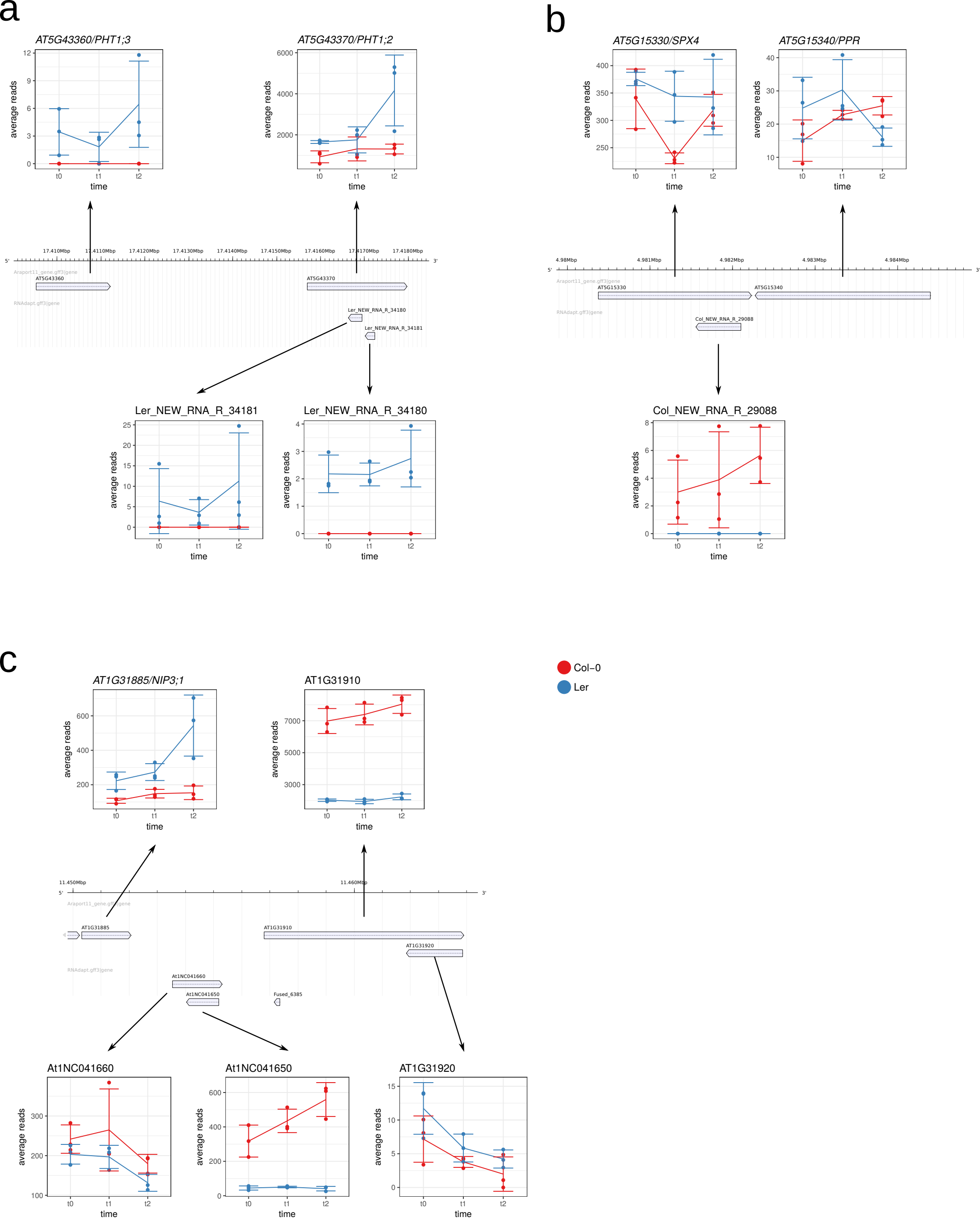

### FigS6.pdf

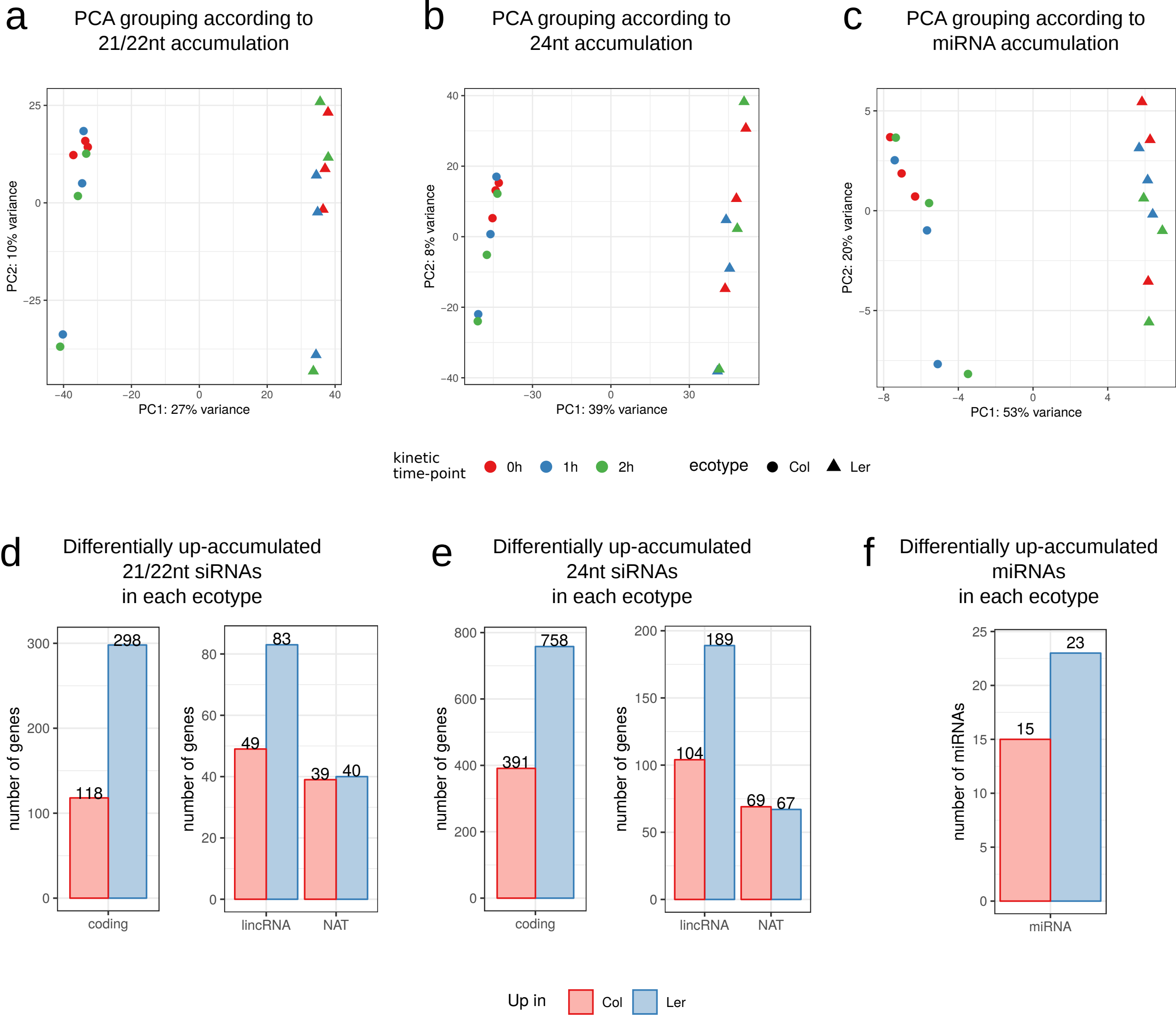

### FigS7.pdf

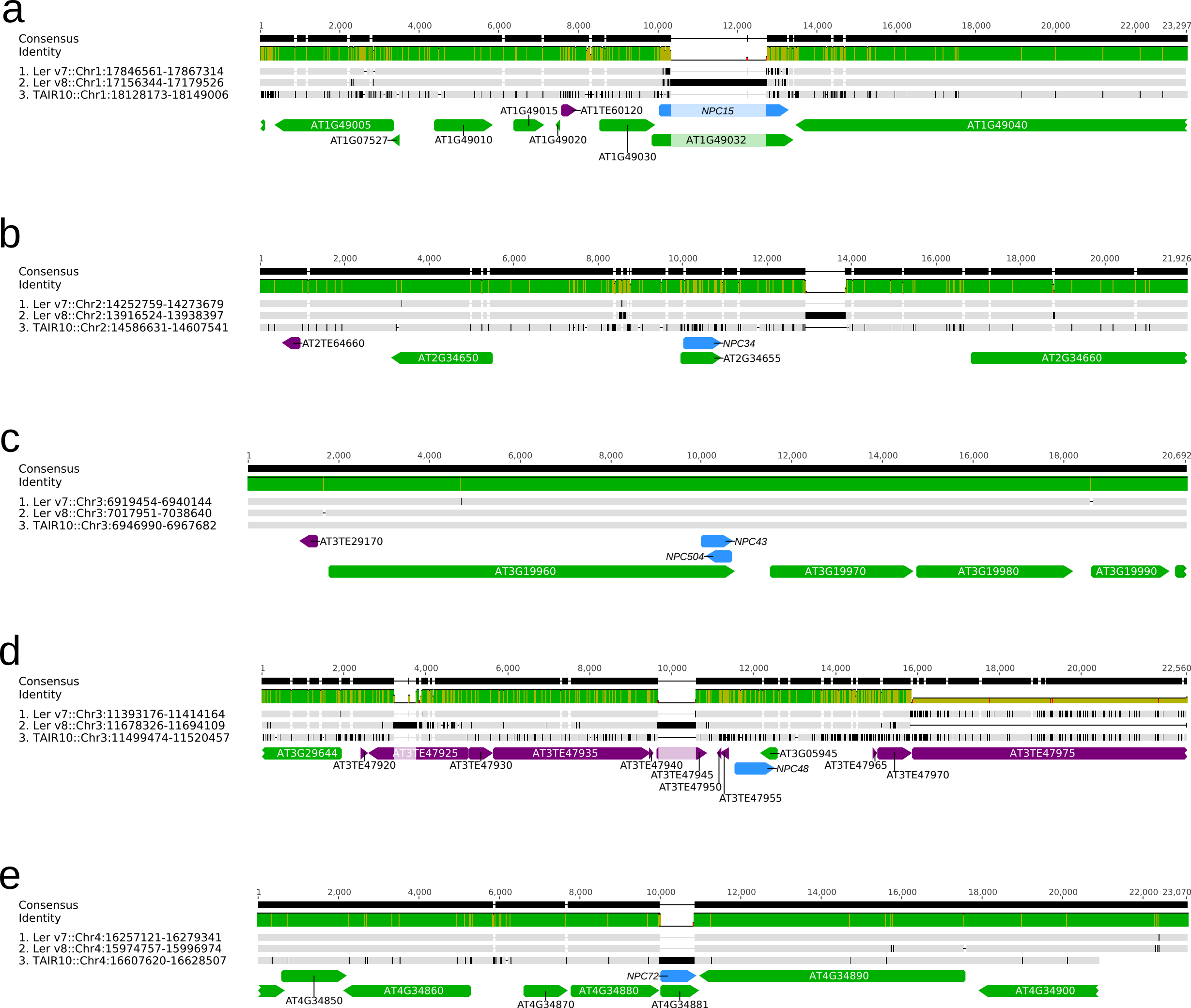

### FigS8.pdf

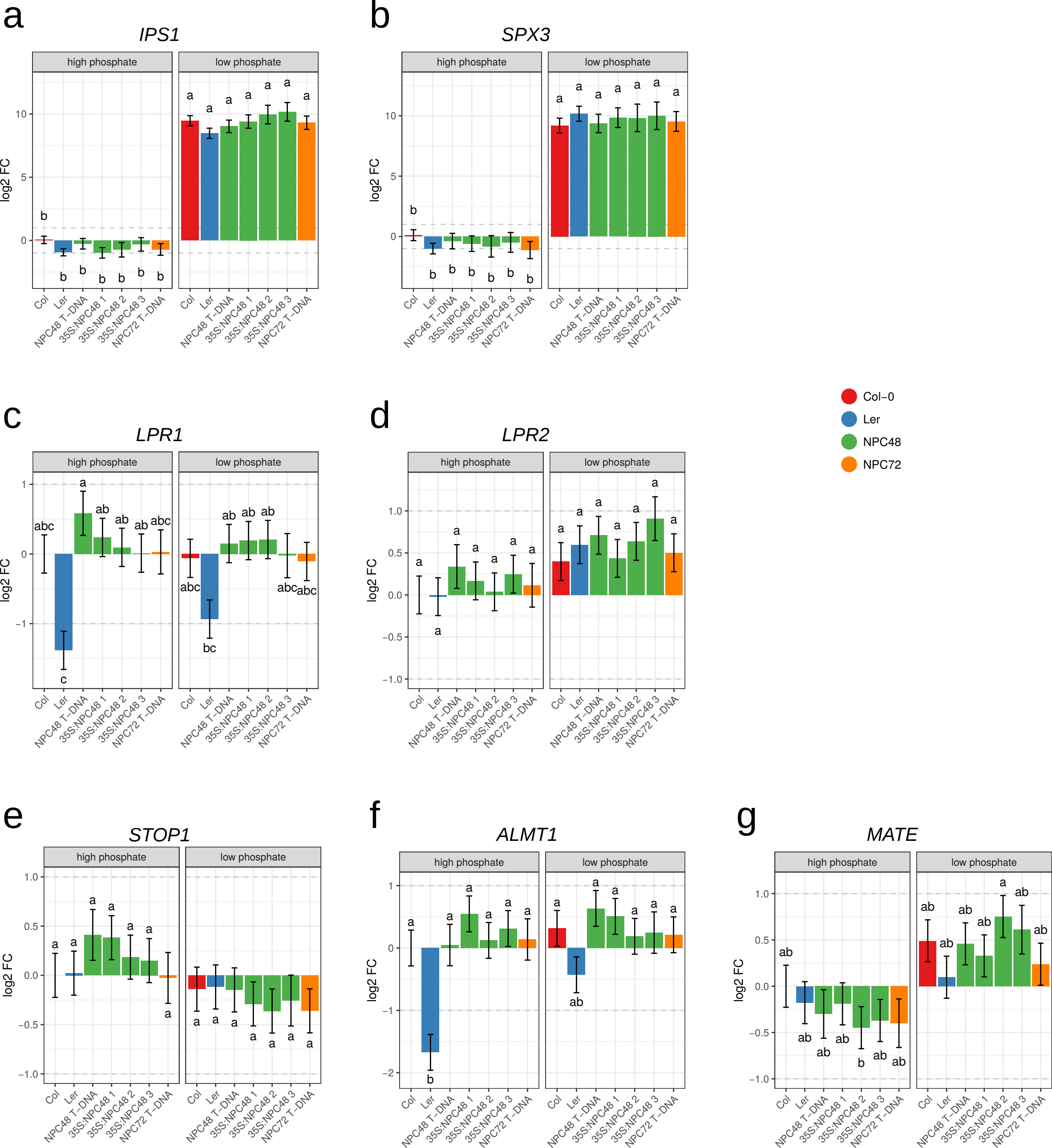
